# Probing Interactions of Therapeutic Antibodies with Serum via Second Virial Coefficient Measurements

**DOI:** 10.1101/2021.06.01.446652

**Authors:** Hayli A. Larsen, William M. Atkins, Abhinav Nath

**Affiliations:** Department of Medicinal Chemistry, University of Washington, Seattle WA 98195, USA

## Abstract

Antibody-based therapeutics are the fastest growing drug class on the market, used to treat aggressive forms of cancer, chronic autoimmune conditions, and numerous other disease states. While the specificity, affinity, and versatility of therapeutic antibodies can provide an advantage over traditional small molecule drugs, their development and optimization can be much more challenging and time-consuming. This is, in part, because the ideal formulation buffer systems used for *in vitro* characterization inadequately reflect the crowded biological environments (serum, endosomal lumen, etc.) that these drugs experience once administered to a patient. Such environments can perturb the binding of antibodies to their antigens and receptors, as well as homo- and hetero-aggregation, in ways that are incompletely understood, thereby altering therapeutic effect and disposition. While excluded volume effects are classically thought to favor binding, weak interactions with co-solutes in crowded conditions can inhibit binding. The second virial coefficient (B_2_) parameter quantifies such weak interactions and can be determined by a variety of techniques in dilute solution, but analogous methods in complex biological fluids are not well established. Here, we demonstrate that fluorescence correlation spectroscopy (FCS) is able to measure diffusive B_2_ values directly in undiluted serum. Apparent second virial coefficient (B_2,app_) measurements of antibodies in serum reveal that changes in the balance between attractive and repulsive interactions can dramatically impact global nonideality. Furthermore, our findings suggest that the common approach of isolating specific components and completing independent cross-term virial coefficient measurements is an incomplete representation of nonideality in serum. The approach presented here could enrich our understanding of the effects of biological environments on proteins in general, and advance the development of therapeutic antibodies and other protein-based therapeutics.

**STATEMENT OF SIGNIFICANCE:** We present FCS as an orthogonal method to traditional methods for characterizing weak, nonspecific homo- and hetero-interactions through determination of self- and cross-term second virial coefficients, respectively. We also characterize weak interactions between therapeutic antibodies and serum components through determination of an apparent second virial coefficient (B_2,app_) directly in undiluted serum. Our results suggest that global nonideality effects are antibody-dependent, and that attractive and repulsive interactions with co-solutes are occurring simultaneously. This approach could advance our understanding of the impact of nonideality to the biophysical and pharmacological properties of therapeutic antibodies and other engineered proteins in relevant biological environments, and could accelerate the development and optimization of protein-based therapeutics.

## INTRODUCTION

Biologics, or protein-based therapeutics, show great promise in treating aggressive cancers, chronic autoimmune conditions, and many other disease states (1). Antibody-based therapeutics, which are a subset of biologics, have emerged as the fastest growing drug class on the market, and often display superior versatility, specificity, and affinity (2). Platforms including conventional monoclonal antibodies (mAbs), bispecific Abs (bsAbs), antibody-drug conjugates (ADCs), Fab fragments, and Fc-fusion proteins have been used successfully in the clinic (3–5). However, new variations of biologics appear in the literature or in pharmaceutical pipelines much more frequently than they reach the clinic or market (6). The inefficient commercial development of biologics reflects the inherent complexity of proteins. Furthermore, our understanding of factors controlling the pharmacokinetics (PK) and disposition of these drugs is extremely limited in comparison to small molecule drugs (7). Biologics interact with their targets and receptors in complex, crowded biological environments such as serum and the endosomal lumen. These biological fluids are not adequately represented in ideal buffer systems used for *in vitro* characterization, which may contribute to the gaps in our knowledge surrounding PK and disposition. One approach to understanding these effects is to investigate how crowded environments affect the biophysical properties of therapeutic antibodies.

Macromolecular crowding is a phenomenon that has been studied extensively and traditionally explained by entropy-driven excluded volume effects (8). Many studies aimed at investigating these effects have utilized various polymers, PEGs, or proteins as crowding molecules to mimic biological environments (9–12). These approaches have many shortcomings, because biological environments are highly concentrated with diverse macromolecules (proteins, lipids, nucleic acids, polysaccharides, etc.). Typical biochemical studies that explore macromolecular crowding are carried out in dilute solutions (1–10 g/L), which are drastically different from biological macromolecular concentrations (50–400 g/L) (13). Excluded volume effects alone enhance stability of proteins by shifting the conformational equilibrium to a more compact state. The equilibrium affinity of proteins for their binding partners may also increase as a result of crowding (9–11). However, these proposed effects do not address the possibility of enthalpically-based weak interactions between proteins and other solutes, which have the potential to negatively impact binding affinity and stability. The complex effects of crowding on therapeutic efficacy and disposition of protein-based therapeutics have not been documented.

The framework of thermodynamic nonideality is often used to describe the weak interactions a protein may encounter in solution (14). Strong interactions are routinely probed using a variety of techniques capable of measuring equilibrium binding constants, while weak interactions are less studied and can be more difficult to probe. The second virial coefficient (B_2_) parameter quantifies such weak interactions and has been used to understand many biophysical properties of proteins in solution such as crystallization, solubility, stability, aggregation, and diffusion (15–20). In general, negative values of second virial coefficients indicate attractive interactions between particles in solution, and positive values indicate repulsive interactions. Self- and cross-term second virial coefficients are used to describe homo- or hetero-interactions, respectively. Methods to determine second virial coefficients include various light scattering techniques (SLS, DLS, CG-MALS) (18, 19), self-interaction chromatography (SIC) (20), membrane osmometry (OM) (21), and analytical ultracentrifugation (AUC) (22). Each of these techniques has its advantages and limitations. For example, DLS is rapid and well-established, but is highly sensitive to aggregates and impurities and is limited to measuring self-term nonideality, while AUC is capable of measuring both self- and cross-term nonideality but is expensive and time consuming.

Because aggregation is a common obstacle in biologics development, many studies have focused on self-term virial coefficients as a predictor of aggregation propensity (20, 23). Cross-interactions of mAbs in solution are not as commonly studied. In this study, we present fluorescence correlation spectroscopy (FCS) as an orthogonal method for calculating both self- and cross-term second virial coefficients. FCS is relatively simple to implement and has the capability of measuring the diffusive properties of molecules in complex media. Recently, AUC measurements have been used to investigate weak interactions between mAbs and serum components through the determination of cross-term virial coefficients for isolated components (24). However, our findings suggest that this reductionist approach may fail to provide a complete representation of the nonideal behavior of mAbs in biological environments. FCS provides an alternative approach to measure an apparent second virial coefficient, B_2,app_, for the “bulk environment” for three different mAbs in fetal bovine serum (FBS). These measurements serve to probe the global nonideality of mAbs in serum, rather than cross-terms between specific components, to better understand the forces acting on therapeutics once introduced into a patient. The advantage of this approach is that it incorporates the net entropic and enthalpic effect of all solution components, rather than the effects of single pair-wise interactions. While this study investigates mAbs in serum, this approach has the potential to be expanded to other antibody platforms, protein systems, and biological fluids. The ability to probe nonideality directly in biological fluids could help reveal the impacts of weak interactions on the biophysical properties of biologics in relevant environments.

## THEORY

The virial coefficients describe deviations from ideal behavior in fluids due to pair-wise or higher-order interactions among the constituent molecules. So, for example, pressure *P* can be expressed as a virial equation in terms of a power series in the number density *ρ* (*n*/*V*):

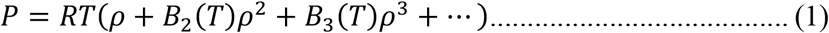

where *R* is the gas constant, *T* is temperature, and *B_n_* is the *n*th virial coefficient which corresponds to interactions between *n* molecules (14). Similarly, the virial coefficients capture deviations from ideality in osmotic pressure Π:

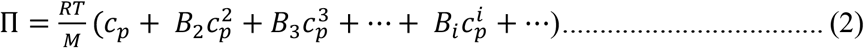

Here, *c_p_* is the mass concentration and *M* is the molecular weight. The sign of the virial coefficient indicates whether pressure is higher or lower than an ideal fluid. For example, if *B_2_* is positive, then the pressure is higher than an ideal fluid due to repulsive pairwise intermolecular interactions between molecules. Conversely, a negative *B_2_* indicates attractive pairwise interactions between molecules. For the remainder of this paper, we neglect the third- and higher-order terms, which are in general much smaller than the second-order term.

Virial coefficients also affect the diffusion of molecules, since repulsive and attractive interactions result in greater and less displacement, respectively, in a given time period. The diffusive properties of molecules can therefore be used to determine second virial coefficients, following the derivation of Harding and Johnston (25). In ideal, dilute solutions, the diffusion coefficient is related to the hydrodynamic radius *R_H_* as expressed by the Stokes-Einstein equation:

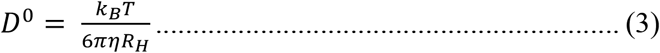

where *k_B_* is Boltzmann’s constant, *T* is absolute temperature, and *η* is viscosity. The diffusion coefficient in the presence of higher concentrations of solute or co-solutes is also related to the concentration gradient of the osmotic pressure, and hence the virial coefficients, as follows:

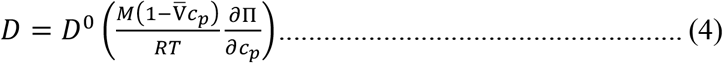

Here, 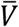 represents the partial specific volume, which is usually assumed to be a constant equal to ~0.7 mL/g for globular proteins. Differentiating eq. (1) with respect to concentration yields:

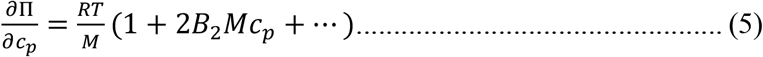

Substituting eq. (5) into eq. (4) and simplifying then results in the following relationship, which is accurate to the first order:

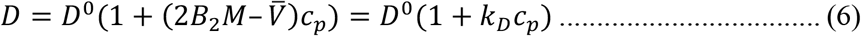

To reiterate, here *D* is the diffusion coefficient observed at a given concentration of solute or co-solutes (corrected for changes in the bulk viscosity of the solution), *D*^0^ is the diffusion coefficient in dilute solution, *M* is molecular weight of unlabeled species, *B*_2_ is the second osmotic virial coefficient, *c_p_* is the concentration of the unlabeled species (g/ml), and *k_D_* is the diffusion interaction parameter defined by:

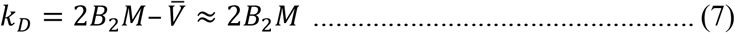

With a complex mixture such as serum, the appropriate value of M (i.e., the effective molecular weight of the various interacting species) is not clear. Reporting results in terms of *k_D_* or 2*B*_2_*M*, rather than *B*_2_, avoids this problem.

Note that *k_D_* is distinct from the equilibrium dissociation constant *k_D_* or the dissociation rate constant *k_d_*. Furthermore, our definition of *k_D_* differs from many earlier works (24, 26, 27) that also include a sedimentation interaction parameter *k_s_*. Our correction for changes in bulk solvent viscosity, described in the FCS section below, removes the need to include or estimate this parameter explicitly. This is advantageous because *k_s_* determination typically requires time-consuming and expensive SV-AUC experiments. We and others (28–30) therefore report *k_D_* values, which are proportional to 2*B*_2_*M* and can conveniently quantify thermodynamic nonideality.

As can be seen from eq. (6), when the diffusion coefficient is plotted against solute or co-solute concentration, the slope of the line divided by the *y*-intercept (*D*^0^) yields *k_D_*. FCS and DLS are very similar in that they both measure changes in intensity over time to determine translational diffusion coefficients. DLS monitors changes in the intensity of scattered light while FCS monitors changes in fluorescence intensity, but determination of the second virial coefficient from measured diffusion coefficients is essentially the same. Depending on whether the molecule of interest is monitored in the presence of increasing concentrations of itself or of other co-solutes, the diffusion interaction parameter represents self-term nonideality (denoted as k_22_ or B_22_), or cross-term nonideality (denoted as k_12_ or B_12_) respectively to represent intermolecular interactions between two of the same molecules or two different types of molecules, respectively. When measuring k_12_, intermolecular interactions between two labeled molecules (denoted as k_11_) are considered negligible due to the low concentration of tracer (<100 nM).

### Dynamic light scattering (DLS)

Particles in solution are in Brownian motion and this constant, random motion causes the intensity of scattered light to fluctuate as a function of time. For a single diffusing species, DLS intensity time traces can be fit to the following correlation function (31):

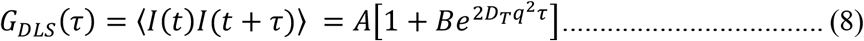

where *τ* is the autocorrelation lag time of the correlator, *q*^2^ is the Bragg wave vector, *D* is the translational diffusion constant, and *A* and *B* respectively represent the baseline and intercept of the correlation function. For a monodisperse system, *q*^2^ depends on the solvent refractive index *n*, wavelength of incident light *λ*, and scattering angle *θ* as follows:

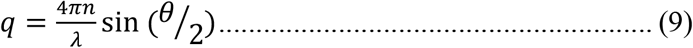

However for polydisperse samples, a variety of analytical approaches have been developed to account for unimodal or multimodal particle size distributions (31). Using such approaches, it is possible to estimate the diffusion coefficient of the species of interest.

### Fluorescence correlation spectroscopy (FCS)

FCS measures the diffusive properties of low concentrations of fluorescent molecules as they move through a well-defined, confocal detection volume. FCS relies on understanding how the fluorescence intensity fluctuates over time. From an intensity time trace, an autocorrelation is calculated and may be fit to yield the diffusion time, *τ_D_* using the following equation (32):

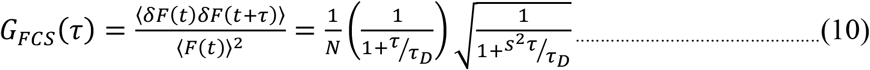

where N is the mean number of molecules in observation volume, *τ_D_* is the correlation decay time due to translational diffusion, and *s* is the axial ratio of the detection volume (0.2 for our instrument). The diffusion time obtained from fitting FCS traces relate to the diffusion coefficients as follows:

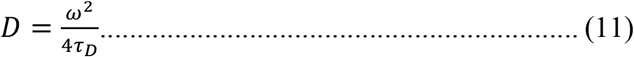

where *ω* is the radius of the confocal volume in the *x*-dimension. In this study, ω^2^ was experimentally determined based on the diffusion time of free probe (Alexa Fluor 488) with known Stokes radius in buffer, using the following equation:

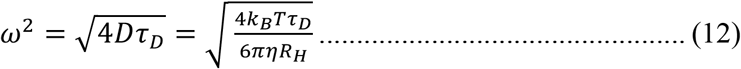

In order to measure *k_D_* using a D vs. c plot (eq. 6), it is necessary to correct for changes in bulk solution viscosity, so that the observed dependence of diffusion time of labeled protein (at a constant, low nM concentration) depends solely on the second virial interactions with unlabeled carrier proteins (the identical protein for B_22_ measurements, a different protein for B_12_ measurements, or a complex mixture such as serum for B_2,app_ measurements). Therefore, the viscosity of carrier protein stocks were measured by FCS, using passivated Alexa Fluor 488 (R_H_ = 5.80 x 10^−8^ cm (33)) as a standard. The viscosity of intermediate concentrations of carrier protein were calculated by linear interpolation based on the stock dilution. Diffusion times were then scaled by the fold change in viscosity:

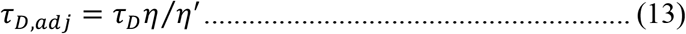

where *τ_D_* is the observed diffusion time, η is the buffer viscosity, and η’ is the viscosity of the carrier protein solution. These adjusted diffusion times measured over a range of carrier protein concentrations were used to calculate diffusion coefficients using eq. (11), which were then fit to eq. (6) to yield a *k_D_* (or 2*B*_2_*M*) value.

## MATERIALS AND METHODS

### Samples

Bovine serum albumin (factor V and protease free) was purchased as a lyophilized powder from GoldBio (St. Louis, MO). Recombinant human serum albumin (HSA) was purchased from Albumin Bioscience (Huntsville, AL). The NIST mAb humanized IgG1 antibody (10 mg/mL) was purchased from the National Institutes of Standards and Technology (RM 8617). Tocilizumab (35 mg/mL) and Carlumab (4 mg/mL) IgG antibodies were provided by the Genentech Outgoing Materials Transfer Agreement program and by Janssen Pharmaceuticals, respectively. FCS and DLS experiments were run in phosphate-buffered saline (PBS: 150 mM NaCl, 2.7 mM KCl, 10 mM Na_2_HPO_4_, and 1.8 mM KH_2_PO_4_, pH 7.4 or 6.0). The antibodies and HSA were stored at 4°C in PBS pH 7.4, while BSA was stored in PBS pH 7.4 or 6.0 depending on the experiment. Fetal bovine serum (FBS) was purchased from Thermo Scientific (A3160401) and was stored in 1 mL aliquots at −20°C until use.

### Protein Labeling

Following the Thermo Scientific labeling protocol, the NIST mAb, tocilizumab, carlumab, and BSA were labeled with Alexa-Fluor 488 carboxylic acid, succinimidyl ester (Thermo Scientific). Desalting was carried out using Zeba Spin Desalting Columns (Thermo Scientific), following the manufacturer’s protocol. Typically, 2-3 Alexa-488 molecules covalently attached to each mAb, while 1-3 Alexa-488 molecules covalently attached to BSA. Labeling efficiency was determined through UV-Vis spectroscopy (A280 and A494 measurements) on a Nanodrop One Microvolume Uv-Vis spectrometer (Thermo Scientific, ND-ONE-W) as well as through diffusion time (change from free dye to protein-bound dye). A488-mAbs and A488-BSA were stored in 1x PBS pH 7.4 at 4°C.

### FCS

All experiments were carried out at room temperature on a home-built instrument based on a Zeiss Axio Observer D1 microscope equipped with Hydra-Harp 400 detection electronics, Tau-SPAD photon counting detector, and pulsed 485 nm laser line driven by a PicoQuant PDL 828 Sepia II driver (PicoQuant GmbH, Berlin, Germany). Sample aliquots of 50 μL were placed on a 22×22 cover glass (VWR 48366-067). Five 30 s measurements of 10 nM Alexa Fluor 488 were used to calibrate the instrument at the start of each experiment. The average diffusion time was used to determine the ω^2^ parameter needed for second virial coefficient calculations. Antibody and BSA measurements (n=5) were carried out for 60s each.

A488-labeled antibodies or BSA were diluted to ~20 nM in varying concentrations of carrier protein (FBS, HSA, or BSA), ranging from 0% to 100%, where the 100% condition ranged from 36–42 mg/mL depending on the experiment.

FCS traces were imported into Prism Graphpad software and fit to a single component FCS equation to yield the average diffusion times at each carrier concentration. These diffusion times were then used to calculate the translational diffusion coefficients at each carrier concentration. Diffusion coefficients *D* were corrected for changes in viscosity and plotted against carrier protein concentration *c*. *D* vs. *c* plots were fit to eq. (6) to determine *k_D_* and *D*^0^. Standard deviations were calculated based on three independent experiments with fresh samples.

The density and viscosity of PBS were measured using an Anton Paar (DMA50000M) densitometer and Anton Paar Automated micro viscometer (AMVn). The viscosity of carrier protein stocks (BSA and FBS) was measured by FCS, using passivated Alexa Fluor 488 as a standard. The starting Alexa Fluor 488 stock was diluted 1000-fold in 150 mM Tris HCl pH 7.4. Passivated Alexa Fluor 488 was further diluted to 10nM in PBS and 100% carrier protein stock solutions. The diffusion time of Alexa Fluor 488 in PBS was used to determine ω^2^ from the previously determined viscosity value using eq. (12) as described in the above Theory section. The *ω*^2^ value, which does not significantly change over the relevant concentration ranges of serum (34), and the average diffusion time from 5 measurements, were used to calculate the diffusion coefficients of the 100% FBS and BSA solutions using eq. (11). The viscosity values were then calculated by rearrangement of the Stokes-Einstein equation and substitution of the determined diffusion coefficients at each condition (Table S2)

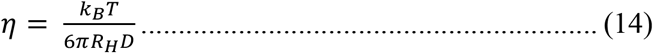

Intermediate viscosity values were obtained from fitting the 0% and 100% viscosity values to a straight line. HSA viscosity values were estimated from BSA values.

### DLS

All dynamic light scattering (DLS) experiments were carried out on a DynaPro Nanostar analyzer (Wyatt Technology) equipped with Dynamics V7 software. BSA samples (100 μL) were spun at 14,000 RPM for 25 minutes at 25°C on an Eppendorf 5415C centrifuge (Brinkmann Instrument Inc) to avoid dust in samples. A quartz cuvette was loaded with ~10 μL of sample directly from the centrifuge tube. Diffusion coefficient measurements (n=20) were carried out at 25°C with a 5 s acquisition time. Viscosity corrections were carried out automatically in the DLS software by adding predetermined viscosity values into the sample parameters for each sample. Data were fit using the regularization analysis in Dynamics V7 (Wyatt Technology) software to generate an intensity versus diffusion coefficient histogram. The average diffusion coefficient of the most intense peak (which represents the monomeric species) was used to calculate *k_D_* or *B*_22_*M*. Using the average diffusion coefficient of the complete distribution had only a minor effect on the *k_D_* values (Fig. S1). Experiments were repeated three times, and averaged *D* values were plotted against carrier concentration and fit to eq. (6) to determine *k_D_* and *D*^0^ values ± SD over three repeats.

## RESULTS

### Validation of FCS-based Virial Coefficient Measurements

In order to validate the use of FCS for the measurement of virial coefficient values, we first compared B_22_ values (i.e., the self-interaction term) for the model protein BSA at pH 7.4 and 6.0 obtained by FCS and by a widely-used dynamic light scattering (DLS) approach. BSA has been extensively studied, and B_22_ values have been reported over a range of different ionic strength and pH conditions. FCS and DLS are related techniques that involve the autocorrelation of a signal from particles in solution to infer their hydrodynamic properties. The key difference is that DLS relies on light scattered by the sample, while FCS relies on fluorescence emitted by the sample. The two techniques each have their advantages and drawbacks: DLS does not require the sample to be labeled but cannot be used to monitor a particular species of interest in a complex mixture. On the other hand, FCS does require the species of interest to be labeled (generally at quite low concentrations, ≤ 100 nM), but can be used with much more challenging samples to answer questions about complex mixtures that are intractable by DLS.

Because FCS and DLS are related, we could employ parallel approaches to obtain B_22_ values by each method as described in the Theory section. FCS and DLS measurements were carried out over a range of concentrations of BSA from 0.38 mg/mL to 38 mg/mL. Autocorrelation curves, generated via either technique, were fit to single-component models (Fig. 1A and Fig. 1B), which treat the relevant signal as originating from a single homogenous species, to obtain diffusion coefficients of BSA at each concentration. Residual free dye in the sample can result in a second, faster-diffusing component in FCS measurements, but based on the calculated labeling efficiency and model comparison calculations (see supporting methods), this does not appear to be a concern for our samples. Another potential concern is the formation of dimers or small soluble oligomers, which can bias the diffusion times to higher values. While accounting for such species using diffusion times alone can be challenging, FCS also yields a brightness per particle parameter that can be obtained by dividing the average intensity by the average particle count (*N* from eq. 10) for a given measurement. Brightness per particle did not vary systematically as a function of carrier protein concentration, and our diffusion coefficients are very similar to those expected of monomeric BSA or antibodies, together suggesting that dimerization or oligomerization do not impact our finding (Table S3). As described in the Theory section, B_22_ values can be calculated by fitting to equation (6). Note that the diffusion coefficients are scaled based on the bulk viscosity of the solution at each concentration of BSA based on predetermined viscosity measurements as described in the Theory section.

**FIGURE 1.**
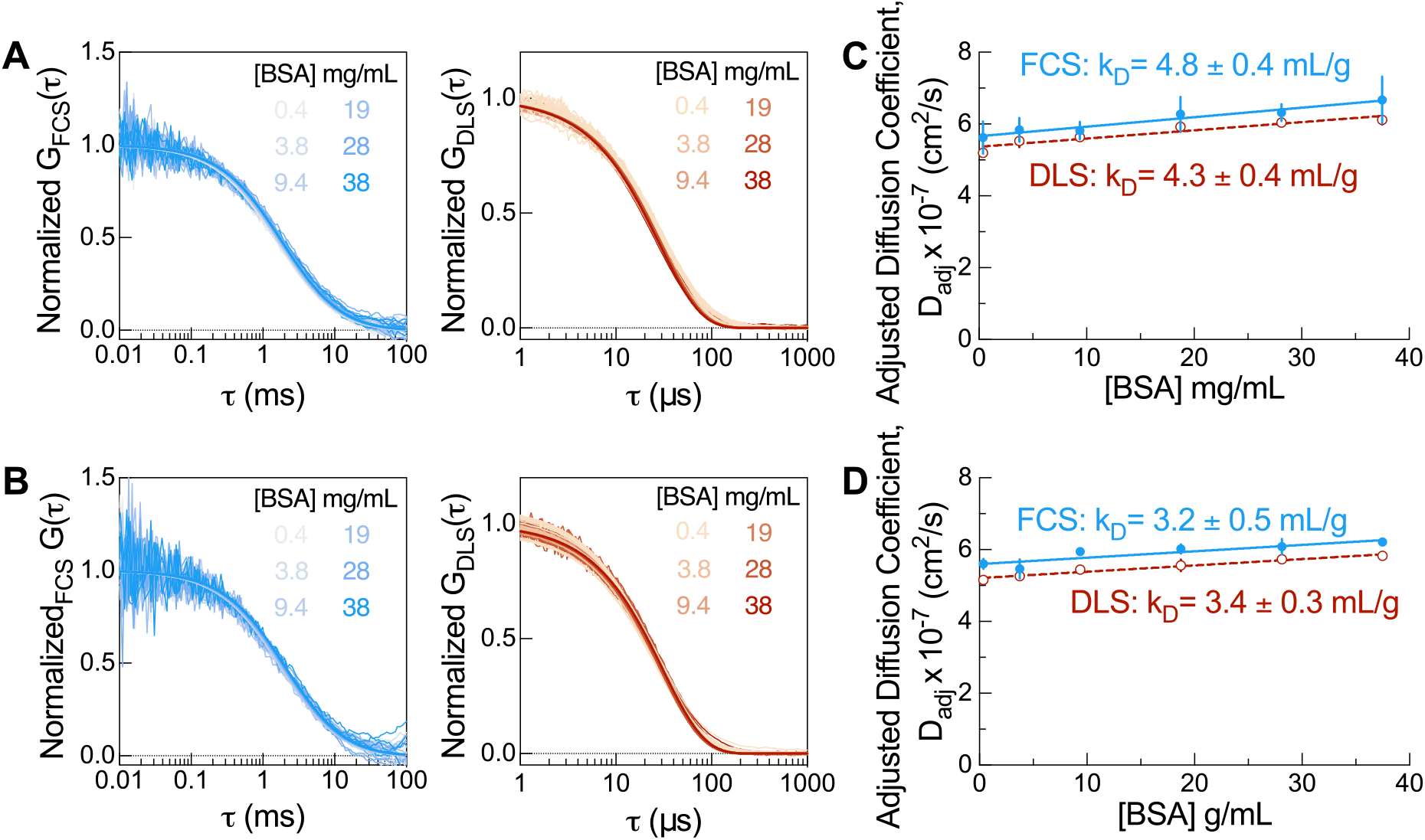
Self-term nonideality for BSA at pH 7.4 and 6.0 measured by FCS and DLS. FCS (left) and DLS (right) traces were collected over a range of BSA concentrations at pH 7.4 (A) and pH 6.0 (B). Diffusion coefficients were scaled based on sample viscosity and plotted against BSA concentration for pH 7.4 (C) and pH 6.0 (D) datasets. These plots were fit to the linear equation *D* = *D*^0^(1 + *k_D_c*) to obtain *k_D_* values.

Fig. 1C shows FCS and DLS comparison of diffusion coefficients for BSA at pH 7.4 out to 37.5 mg/mL. The second virial coefficient, *B*_22_*M* (represented by the interaction parameter *k_D_*) and *D*^0^ values obtained by FCS (4.8 ± 0.4 mL/g and 5.66 ± 0.26 x 10^−7^ cm^2^/s respectively) were comparable to values obtained by DLS (4.3 ± 0.4 mL/g and 5.37 ± 0.09 x 10^−7^ cm^2^/s respectively). The corresponding *B*_22_*M* and *D*^0^ values obtained at pH 6.0 (Fig. 1D) by FCS (3.2 ± 0.5 mL/g and 5.60 ± 0.07 x 10^−7^ cm^2^/s respectively) were also comparable to DLS values (3.4 ± 0.3 mL/g and 5.21 ± 0.04 x 10^−7^ cm^2^/s respectively). These measurements are highly sensitive to buffer conditions resulting in a wide range of second virial coefficient values in the literature, but our results are well within the range of previously reported values (35, 36). The lower *B*_22_ value at pH 6.0 is also consistent with trends in the literature. The positive virial coefficients in both cases indicate weak repulsive interactions between BSA molecules. These results suggest that FCS can be used as an orthogonal method for calculating self-term nonideality.

### NIST mAb Cross-Term Interactions with Albumin

NIST mAb with human serum albumin (HSA) as carrier protein was used as the model system to validate cross-term nonideality measurements by FCS. Fig. 2A shows diffusion coefficients for NIST mAb in solutions containing between 0 and 36 mg/mL HSA. Second virial coefficient *B*_12_*M*, and *D*^0^ values obtained by FCS (3.3 ± 0.2 mL/g and 4.08 ± 0.10 x 10^−7^ cm^2^/s respectively) were comparable to AUC values (3.2 ± 0.3 mL/g and 4.17 ± 0.03 x 10^−7^ cm^2^/s) reported by Wright et al. (24). This result suggests that FCS can be used as an alternative method to calculate cross-term nonideality in addition to self-term nonideality. The positive virial coefficient indicates repulsive interactions between NIST mAb and HSA. Cross-interactions between NIST mAb and bovine serum albumin (BSA) were also measured, for comparison with subsequent measurements in fetal bovine serum (FBS). Fig. 2B shows diffusion coefficients for NIST mAb in solutions containing between 0 and 42 mg/mL BSA. At 5.8 ± 1.1 mL/g, the *k_D_* value with BSA was slightly higher than with HSA, while the *D*^0^ value was similar at 4.10 ± 0.12 x 10^−7^ cm^2^/s. The increased *k_D_* suggests stronger repulsive interactions between NIST mAb and BSA than for HSA.

**FIGURE 2.**
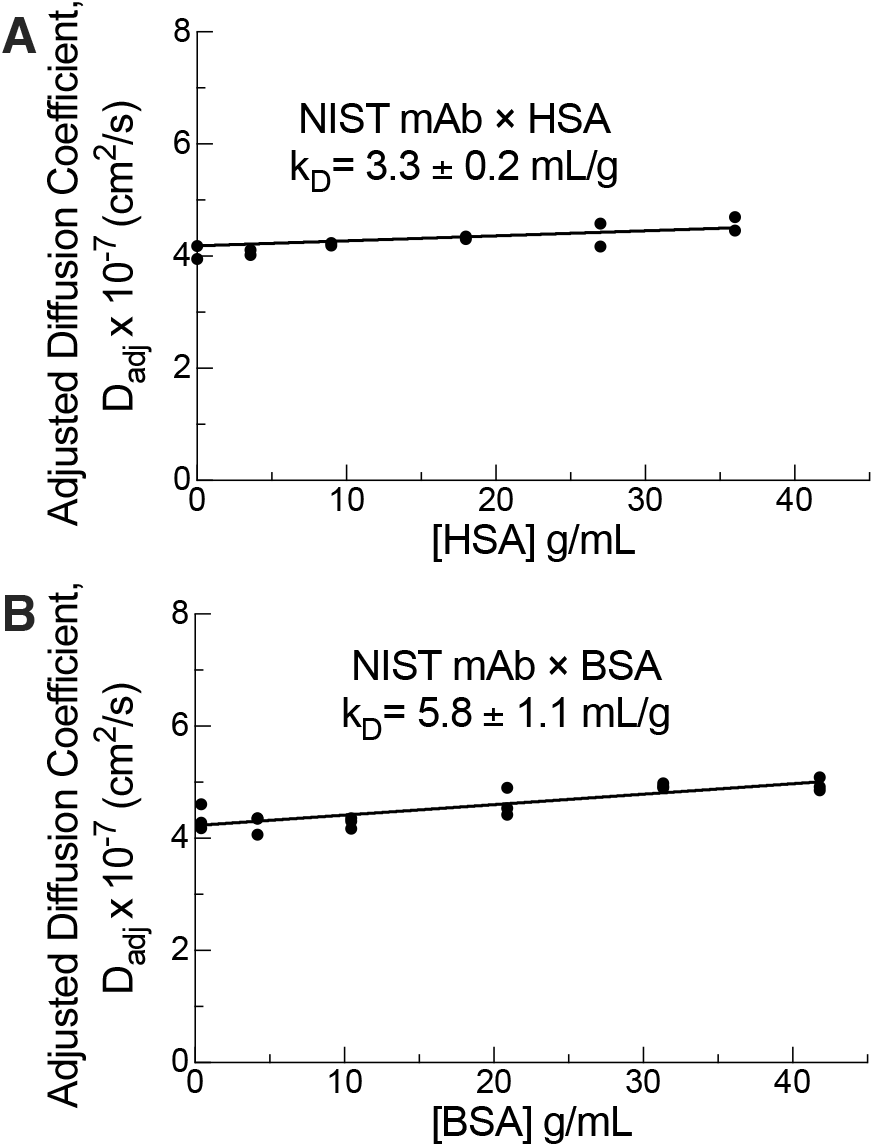
Cross-term nonideality of A488-NIST mAb and human serum albumin (A) or bovine serum albumin (B). Data were fit to the linear equation *D* = *D*^0^(1 + *k_D_c*) to obtain *k_D_* values. Both conditions yield positive *k_D_* values, indicating repulsive interactions between NIST mAb and albumin.

Albumin is the most abundant protein in serum, making it a good carrier protein to investigate cross-term nonideality for protein-based therapeutics (37). The repulsive interactions observed between NIST mAb and HSA could shift the folding equilibrium of the antibody to favor more compact structures, thus enhancing folding stability. Further assessment needs to be completed with a broader panel of antibodies to determine potential trends; however, these effects could be antibody dependent. Despite the abundance of albumin in serum, there are many other components to consider that could have opposing effects. Therefore, cross-term measurements with albumin alone may be an incomplete representation of the nonideality exhibited by a protein-based therapeutic in serum.

### NIST mAb, Tocilizumab, and Carlumab Cross-term Interactions with Serum

We introduce an apparent second virial coefficient, B_2,app_, which measures global nonideality between a labeled species and serum components. By labeling antibodies and diluting them in varying concentrations of serum, we were able to probe nonideality in complex media. In this approach, we are unable to specify which component(s) in serum are interacting with our labeled antibodies. It is worth noting that a *k_D_* value of 0 mL/g does not necessarily indicate a lack of interactions with serum components, but could instead reflect a balance of both attractive and repulsive interactions. The positive and negative values resulting from repulsive interactions with one component in the medium and attractive interactions with another component would cancel each other out. A value deviating from 0 mL/g would indicate that either repulsive or attractive interactions dominate but does not exclude the possibility of opposing interactions occurring simultaneously. In theory, B_2,app_ could also be applied to other complex multicomponent systems such as plasma, endosomal lysate, etc.

B_2,app_ values in serum were determined for three antibodies: NIST mAb, Carlumab, and Tocilizumab. NIST mAb served as a reference mAb, while Tocilizumab is a clinically approved therapeutic (an interleukin-6 receptor inhibitor used to treat autoimmune diseases) and Carlumab is a discontinued therapeutic candidate. Measurements of A488-NIST mAb, A488-Tocilizumab and A488-Carlumab with fetal bovine serum (FBS) as the carrier solution were used to calculate apparent second virial coefficients as shown in Fig. 3. FCS traces (not shown) were fit to a single-component FCS equation to yield diffusion time (*τ_D_*) values at each FBS concentration. Adjusted diffusion coefficients, *D_adj_*, were calculated from diffusion times that were adjusted based on separately determined solution viscosity data. *D_adj_* values were plotted against FBS concentration and data were fit to yield *k_D_* or 2*B_2,app_M* values. A488-NIST mAb yielded a *k_D_* value of −0.2 ± 0.3 mL/g and a *D*^0^ value of 4.04 ± 0.12 x 10^−7^ cm^2^/s (Fig. 3A). A488-Tocilizumab yielded a similar result with *k_D_* value of 0.2 ± 0.5 mL/g and a *D*^0^ value of 4.42 ± 0.26 x 10^−7^ cm^2^/s (Fig. 3B). In both cases, *k_D_* values did not significantly deviate from 0 mL/g. A488-Carlumb exhibited a *k_D_* value of −10.4 ± 0.9 mL/g and a *D*^0^ value of 4.56 ± 0.14 x 10^−7^ cm^2^/s (Fig. 3C), suggesting weak attractive interactions with serum components. The possible curvature and the magnitude of the change observed in the *D_adj_* vs. concentration plot of A488-Carlumab vs. FBS could indicate the presence of more complex or higher-order interactions with serum components, but this needs further investigation. Interestingly, the discontinued therapeutic displays markedly different behavior than the other two mAbs, with (on balance) much stronger attractive interactions with serum components.

**FIGURE 3.**
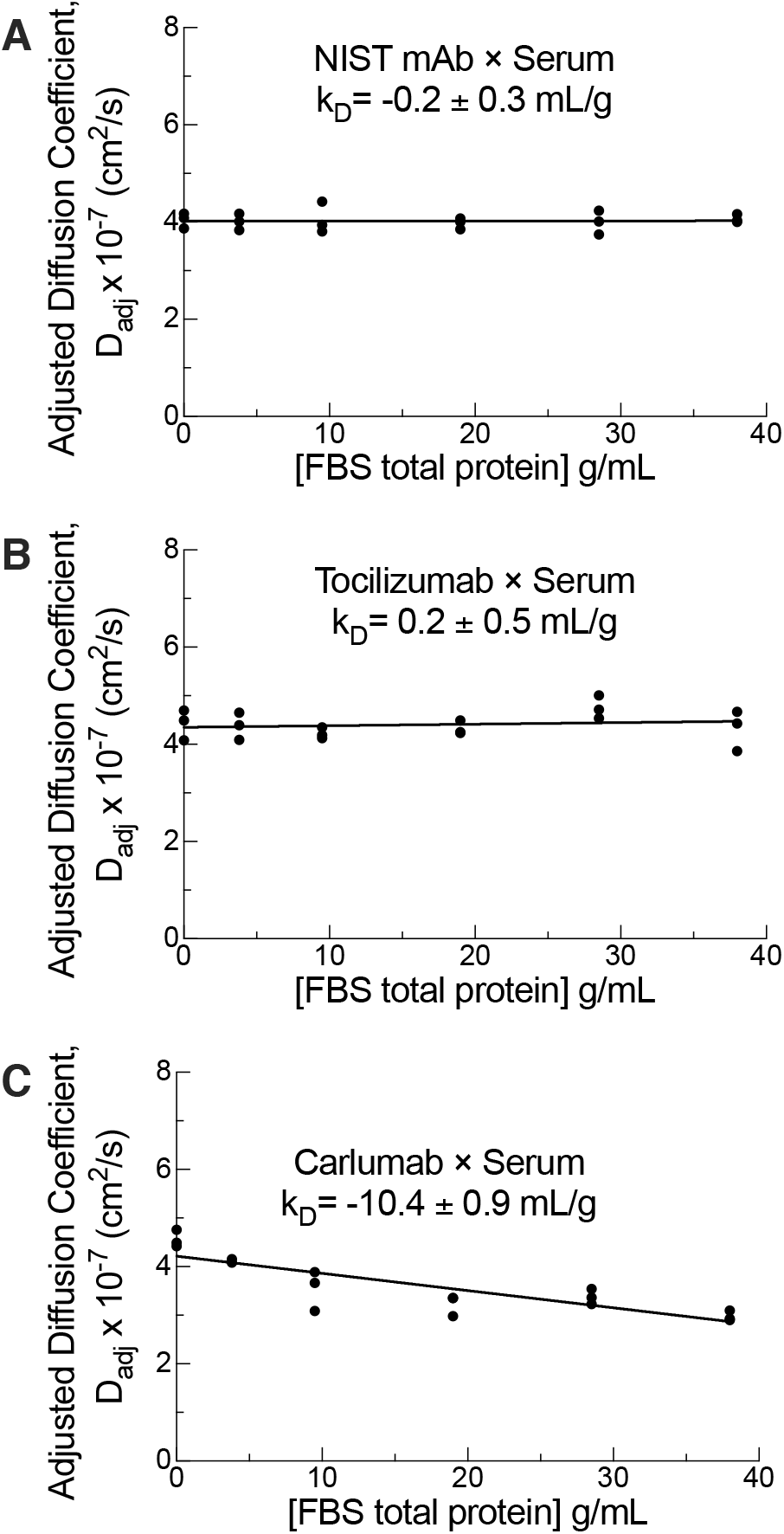
Apparent second virial coefficients for mAbs in FBS: D_adj_ vs. [FBS] for NIST mAb (A), Tocilizumab (B), and Carlumab (C). NIST mAb and Tocilizumab show negligible deviations from ideal behavior, while Carlumab shows a marked decrease in diffusion coefficient indicative of attractive interactions with one or more co-solutes in serum.

Table 1 summarizes both self and cross-term nonideality parameters obtained from DLS and FCS measurements. Validation of our FCS method was accomplished with BSA and NIST mAb model systems. Determination of the diffusion interaction parameter *k_D_* for BSA at pH 7.4 and 6.0 yielded comparable results between FCS and DLS methods. Similarly, NIST mAb and BSA, our model system for cross-term nonideality, yielded results comparable to the AUC literature value. We introduced an apparent second virial coefficient as a determinant of global nonideality in complex media and successfully measured *k_D_* values for three mAbs (NIST mAb, Tocilizumab, and Carlumab) in FBS.

**Table 1.**
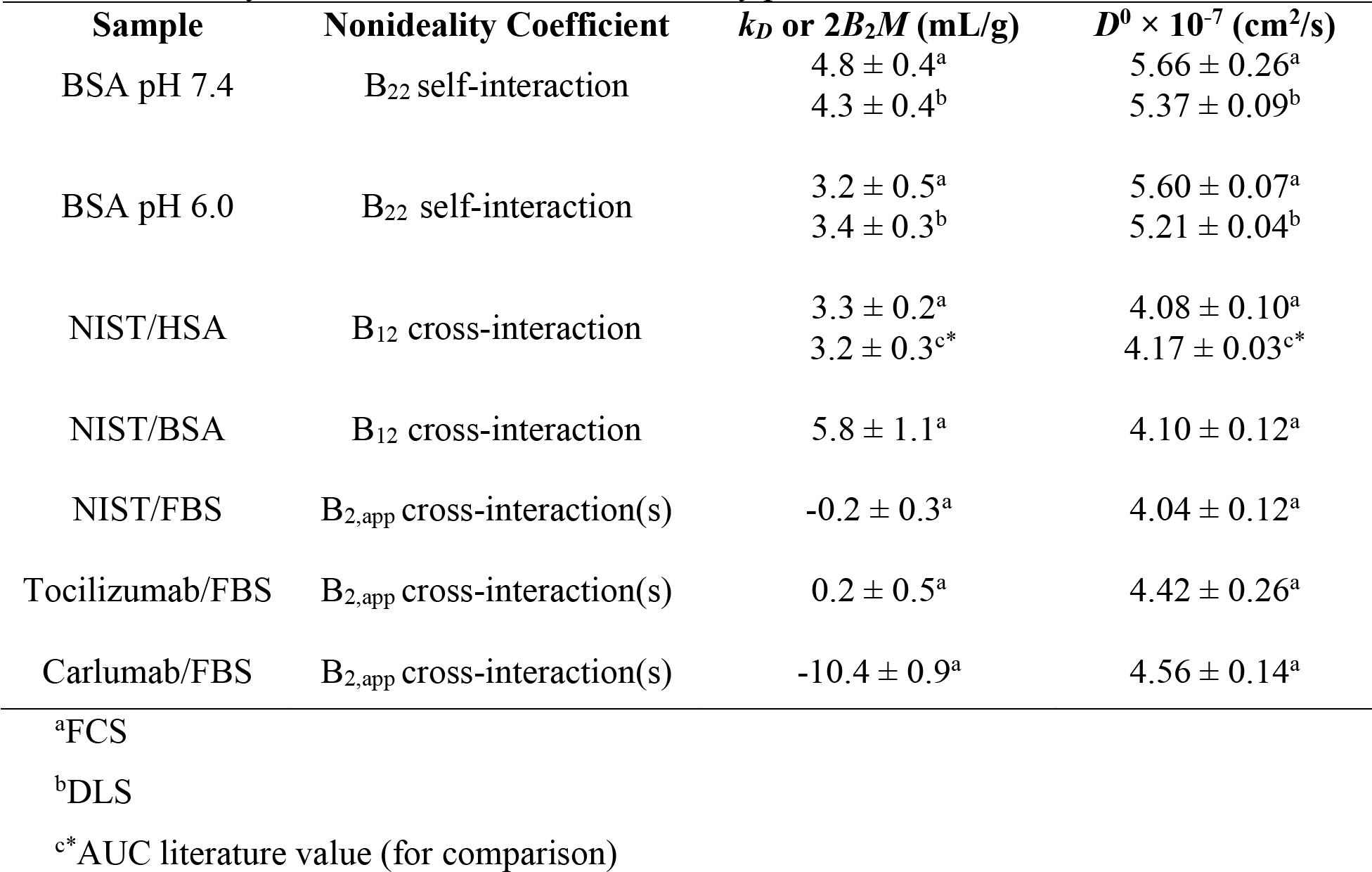
Summary of self- and cross-term nonideality parameters.

| Sample | Nonideality Coefficient | $k_D$ or $2B_2M$ (mL/g) | $D^0 \times 10^{-7}$ (cm<sup>2</sup>/s) |
| --- | --- | --- | --- |
| BSA pH 7.4 | B<sub>22</sub> self-interaction | $4.8 \pm 0.4^a$ | $5.66 \pm 0.26^a$ |
| | | $4.3 \pm 0.4^b$ | $5.37 \pm 0.09^b$ |
| BSA pH 6.0 | B<sub>22</sub> self-interaction | $3.2 \pm 0.5^a$ | $5.60 \pm 0.07^a$ |
| | | $3.4 \pm 0.3^b$ | $5.21 \pm 0.04^b$ |
| NIST/HSA | B<sub>12</sub> cross-interaction | $3.3 \pm 0.2^a$ | $4.08 \pm 0.10^a$ |
| | | $3.2 \pm 0.3^{c*}$ | $4.17 \pm 0.03^{c*}$ |
| NIST/BSA | B<sub>12</sub> cross-interaction | $5.8 \pm 1.1^a$ | $4.10 \pm 0.12^a$ |
| NIST/FBS | B<sub>2,app</sub> cross-interaction(s) | $-0.2 \pm 0.3^a$ | $4.04 \pm 0.12^a$ |
| Tocilizumab/FBS | B<sub>2,app</sub> cross-interaction(s) | $0.2 \pm 0.5^a$ | $4.42 \pm 0.26^a$ |
| Carlumab/FBS | B<sub>2,app</sub> cross-interaction(s) | $-10.4 \pm 0.9^a$ | $4.56 \pm 0.14^a$ |

<sup>a</sup>FCS
<sup>b</sup>DLS
<sup>c\*</sup>AUC literature value (for comparison)

## DISCUSSION

Studies of the second virial coefficient date back over a century, and their original application to gases has broadened to encompass many other systems. Although derived from the ideal gas law, similar deviations from ideality apply to diffusion and sedimentation behavior, making it possible to measure second virial coefficients with many analytical techniques. The second virial coefficient in protein systems has been extensively studied in terms of the effects of solution conditions, such as pH and ionic strength, as well as of weak protein interactions. Methods used include membrane osmometry, AUC, size-exclusion chromatography (SEC), cross-interaction chromatography (SIC), and various light scattering techniques such as DLS, static light scattering (SLS), and composition-gradient multi-angle light scattering (CG-MALS). There are disadvantages associated with each of these methods. For example, chromatography methods often consume a lot of protein and immobilization conditions can be challenging to establish. Membrane osmometry is complex and requires determination of specific properties for the individual proteins prior to analysis. Light scattering measurements such as DLS are highly sensitive to impurities and, as previously mentioned, cannot measure cross-term nonideality. AUC requires costly instrumentation as well as expertise in running experiments and analyzing sedimentation data. The capability of fluorescence detection in AUC measurements (AU-FDS) allows AUC to be applied to complex media, but the analysis is complicated by the fact that the concentration and viscosity of the medium varies over the sample. Sedimentation and diffusion coefficients are system properties that depend on the concentration of other solutes present in solution. Furthermore, the viscosity of the samples varies depending on position in the sample cell during centrifugation. As a result, AUC requires analysis of both sedimentation and diffusion interaction parameters, *k_S_* and *k_D_* respectively, to determine the second virial coefficient. While second virial coefficient determination has been reported by AUC in buffer systems, determination in complex biological fluids has not been reported. The interpretation of such data is complicated, requiring a more detailed analysis than currently available. The correction for changes in bulk viscosity in our FCS measurements, as discussed in the above Theory section, eliminates the need to estimate the *k_S_* parameter, thus simplifying our analysis. We have successfully validated FCS as an orthogonal method for calculating both self and cross-term virial coefficients. FCS experiments are relatively fast and do not consume large quantities of protein. One major downfall of this technique, as discussed in the Results section, is that FCS does require the species of interest to be labeled. However, A488 labeling of mAbs and BSA in this study was efficient and does not appear to have perturbed interactions with co-solutes. FCS is relatively inexpensive and easy to implement, making it an attractive alternative to other methods.

The second virial coefficient is finding applications in the biopharmaceutical industry, as a tool to better understand protein aggregation. The self-term virial coefficient (*B*_22_) and kD have demonstrated applicability (27, 38–41) in relating intermolecular interactions to various biophysical properties of proteins, including aggregation. Biologics are subject to a range of stresses during manufacturing such as variations in ionic strength, pH, temperature, and high protein concentrations that can drive aggregation. The negative impacts of aggregation are not limited to manufacturing but can also be detrimental to half-life, efficacy, and the safety profiles of therapeutic products (42). As a result, the biopharmaceutical industry has shown increasing interest in utilizing *B*_22_ as a predictor of aggregation propensity. In contrast, measurements of cross-term second virial coefficients (*B*_12_) have been underutilized. Perhaps the most common application of cross-term virial coefficient determination is to probe protein-excipient interactions during formulation development (41). There is, however, an emerging movement towards studying weak interactions in crowded conditions. While macromolecular crowding has traditionally focused on excluded volume effects, there is increasing agreement that weak interactions in crowded conditions have the potential to overcome the (generally stabilizing) excluded volume effect. Thus, our understanding of nonideality has seen a paradigm shift. In recent years there has been increasing interest in exploring cross-interactions between therapeutic antibodies and serum proteins. Wright, et al. (24) proposed a preclinical AUC method to measure weak interactions between mAbs and serum components such as HSA and IgG through determination of second virial coefficients and sedimentation interaction parameters. Kim, et al. (43) used CG-MALS to probe cross-term interactions between mAbs and HSA, but also used BLI to explore the functional consequence of this nonideality on antigen binding. While these studies relied on HSA alone as a model system to determine cross-term nonideality of mAbs in serum, our results caution that such an approach may fail to qualitatively or quantitatively capture the true effects of serum.

Our second virial coefficient results for NIST mAb with BSA (*k_D_* = 5.8 mL/g) and for NIST mAb with serum (*k_D_* = −0.2 mL/g) differ in a particularly interesting way. Because albumin is the most abundant component of serum, one might expect its effects to predominate serum-induced nonideality. Instead, the lower *k_D_* value in FBS suggests that repulsive interactions with BSA and attractive interactions with some other component(s) are occurring simultaneously in serum. Furthermore, this suggests that the approach of isolating specific components and completing independent cross-term virial coefficient measurements is an incomplete representation of nonideality in serum. The ability to directly measure diffusion coefficients in complex media gives FCS an important advantage over existing methods in probing global nonideality in relevant biological fluids. Implementing these measurements into biologics development could prove beneficial on several levels from candidate selection to formulation development. For example, comparing *B*_2,app_ values for a panel of mAbs during candidate selection could facilitate elimination or selection of certain candidates. Additionally, implementing *B*_2,app_ measurements during characterization could serve as a complementary tool to various analytical techniques for assessing the impacts of process changes during recovery, purification, and formulation development. In theory, this approach has the potential to be applied to other biological fluids and protein systems. For example, *B*_2,app_ measurements could be conducted in plasma or endosomal lysate to better understand nonideality following internalization. *B*_2,app_ could be measured in interstitial fluids to model tumor microenvironments or in a solution mocking the serum formulation interface during administration. This approach can potentially be applied to different antibody platforms such as antibody-drug conjugates, Fab fragments, bispecific antibodies, and Fc-fusion proteins, as well many other therapeutic proteins, vaccine platforms and diagnostics.

Serum is a complex solution comprising albumin (roughly two-thirds of the protein content), IgG (~20%), IgA (~4%), IgM (~2%), as well as numerous other lipoproteins, complement factors, transport proteins and smaller osmolytes (44–46). Fig. 4 depicts the increasingly crowded environment that a labeled antibody samples when going from 10% (Fig. 4A) to neat (Fig. 4C) serum, while Fig. 4B represents the intermediate condition of 50% serum. Due to short intermolecular distances in the crowded environment, serum components can experience weak, nonspecific interactions with labeled antibodies and with each other. As previously discussed, repulsive interactions occur between NIST mAb and BSA (Fig. 4D). While our apparent second virial coefficient results cannot identify the various interacting species in serum, they do suggest that these repulsive interactions with BSA are counteracted by attractive interactions with other, non-albumin serum components (Fig. 4E). In the case of Carlumab, attractive interactions with as-yet unidentified serum components dominate, as illustrated in Fig. 4F. It is also possible that Carlumab could be experiencing both weak self-association and association with serum components simultaneously, but further investigation is needed.

**FIGURE 4.**
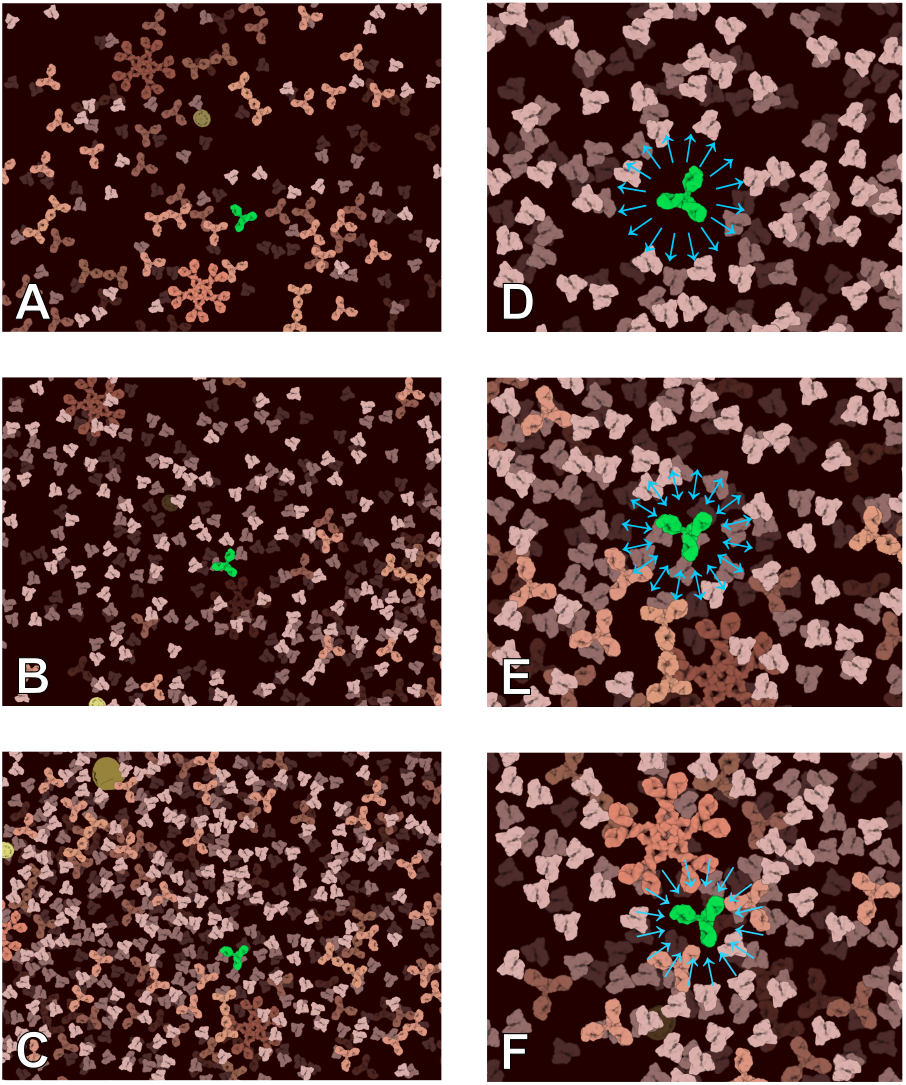
These illustrations, prepared in CellPAINT 2.0 (47), depict the environments experienced by a probe mAb (green) in 10% (A), 50% (B) or 100% serum (C). Co-solutes such as serum albumin, other IgGs, IgAs, IgMs and lipoprotein particles (differing shades of pink) are shown to scale and in approximately the concentrations found in human serum. Molecules of interest may participate in predominantly repulsive interactions with co-solutes, as in the case of NIST mAb and BSA (D). Attractive and repulsive interactions may cancel each other out, as in the case of NIST mAb and serum (E). Finally, attractive interactions may predominate, as in the case of Carlumab and serum (F).

We hypothesize that there may be biophysical consequences of nonideality in crowded biological environments that could impact important properties of biologics (i.e., binding affinity, stability, half-life, etc.). The approach presented here will enable future studies on the effects of serum-induced nonideality on other IgG isotypes, antibody platforms and fragments. Moreover, a component assessment of serum and comparative studies of sera from humans and preclinical species could prove informative. Our comparison of *k_D_* values in BSA and FBS is a preliminary step along this path. Our work also enables mechanistic studies focused on understanding the biophysical basis of nonideality exhibited by mAbs.

## CONCLUSION

In this study, we have validated FCS as an orthogonal method for determining both self and cross-term second virial coefficients via diffusion time measurements and the concentration dependence of corresponding translation diffusion coefficients. Plots of diffusion coefficients against carrier protein concentration were fit to yield diffusion interaction parameter (*k_D_*) values, which are proportional to the second osmotic virial coefficient, 2*B*_2_*M*. Furthermore, the capability of FCS measurements in complex media allowed for determination of an apparent second virial coefficient (*B*_2,app_) for three mAbs in FBS to probe global nonideality effects in serum. These results reveal that multiple forces may be acting on mAbs simultaneously in biological environments, where repulsive or attractive interactions can dominate or balance one another. Further investigation into the biophysical significance of these *B*_2,app_ values is needed. This approach may be useful in areas of biologics development such as candidate selection, characterization, and formulation development, as well as in understanding other biological fluids and protein systems.

## SUPPORTING INFORMATION

### Labeling Efficiency Determination of Alexa488-labeled BSA

The concentration of Alexa488 SE and A488-labeled BSA (A488-BSA) were determined via absorbance measurements to determine the labeling ratio from the following relationships

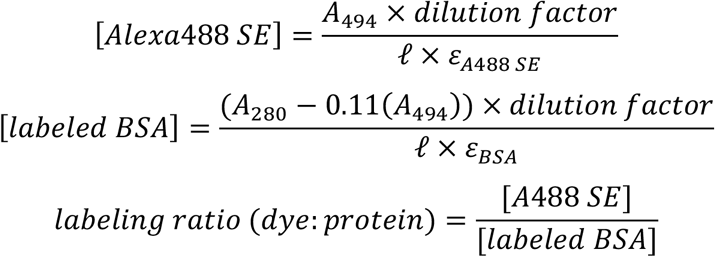

where *A* represents the absorbance at either 280 nm or 494 nm, ℓ represents the path length (1 cm), and *ε* represents the extinction coefficient of either BSA (280 nm) or Alexa488 SE (494 nm). A correction factor of 0.11 is used to account for the contribution of Alexa488 SE to the absorbance at 280 nm based on extinction coefficients provided by the manufacturer (48).

The change in diffusion time between Alexa488 SE and A488-labeled BSA was also used to support labeling efficiency determination by comparing the observed change to the expected fold change in diffusion time. The expected fold change in diffusion time can be estimated as follows

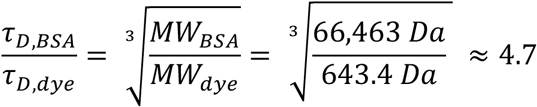

where *Δτ_D_* represents the change in diffusion time and *ΔMW* represents the change in molecular weight, in this case between Alexa488 SE and BSA.

**TABLE S1.**
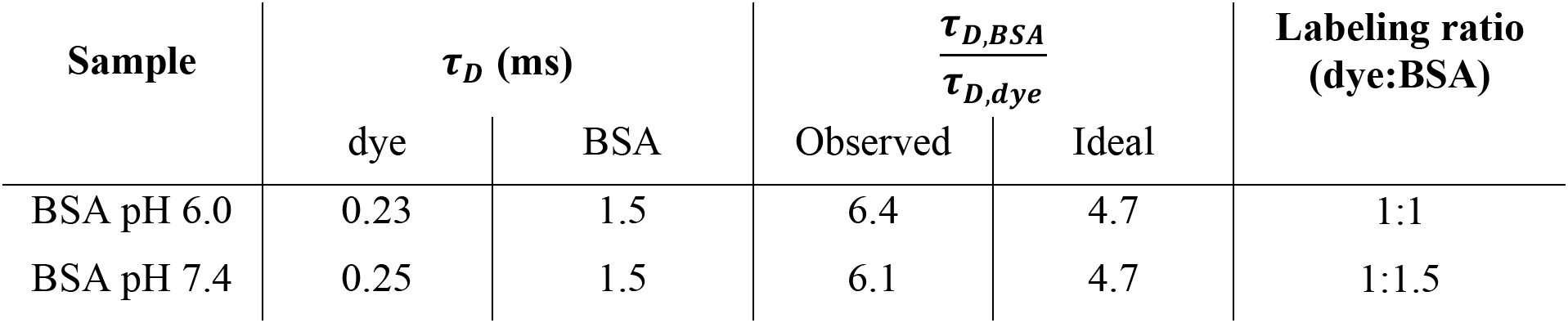
Diffusion time and labeling ratio estimates for BSA model system

### Viscosity Determination of Neat Carrier Protein Solutions

To determine the viscosity of neat carrier protein solutions, the radial dimension of the FCS observation volume (ω^2^ parameter) was first determined from the diffusion time of Alexa488 SE with known Stokes radius (*R_H_*) in 1x phosphate-buffered saline (PBS) pH 7.4 with predetermined viscosity (via densitometer and viscometer measurements as outlined in the Methods section) (*η*) through the following relationship:

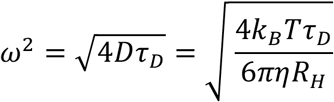

Since the ω^2^ value does not significantly change over the relevant concentration ranges of carrier protein (34), the same value was used to determine the viscosity of neat carrier protein solutions using the Stokes-Einstein equation:

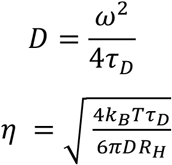

**TABLE S2.** Summary of viscosity values for neat protein solutions

| Sample | Concentration (mg/mL) | Viscosity (cP) |
| --- | --- | --- |
| PBS buffer | N/A | 1.0195 |
| BSA pH 7.4 | 38 | 1.447 |
| BSA pH 6.0 | 38 | 1.280 |
| Fetal bovine serum | 38 | 1.370 |

The viscosity values for lower concentrations of carrier were determined by linear interpolation.

### Brightness per Particle Calculation for BSA

Brightness per particle was calculated by measuring the average intensity <*I*> over the course of an FCS measurement and dividing by the number of molecules, *N*, obtained by least-squares fitting of the autocorrelation to the following equation:

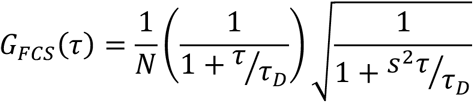

where *τ_D_* is the translational diffusion time and *s* is the axial ratio of the detection volume (fixed to 0.2).

**TABLE S3.** Brightness per particle calculations for BSA model system at pH 7.4 and 6.0.

| [BSA]<br>(mg/mL) | $\langle I \rangle / N$ (Hz) | |
| --- | --- | --- |
|  | pH 7.4 | pH 6.0 |
| 0.38 | 2676 | 1242 |
| 3.8 | 2527 | 1512 |
| 9.4 | 2437 | 1565 |
| 18.8 | 2230 | 1170 |
| 28 | 2257 | 1222 |
| 38 | 2156 | 1263 |

### DLS Virial Coefficient Measurements for BSA Model System from Complete Diffusion Coefficient Distributions

**FIGURE S1.**
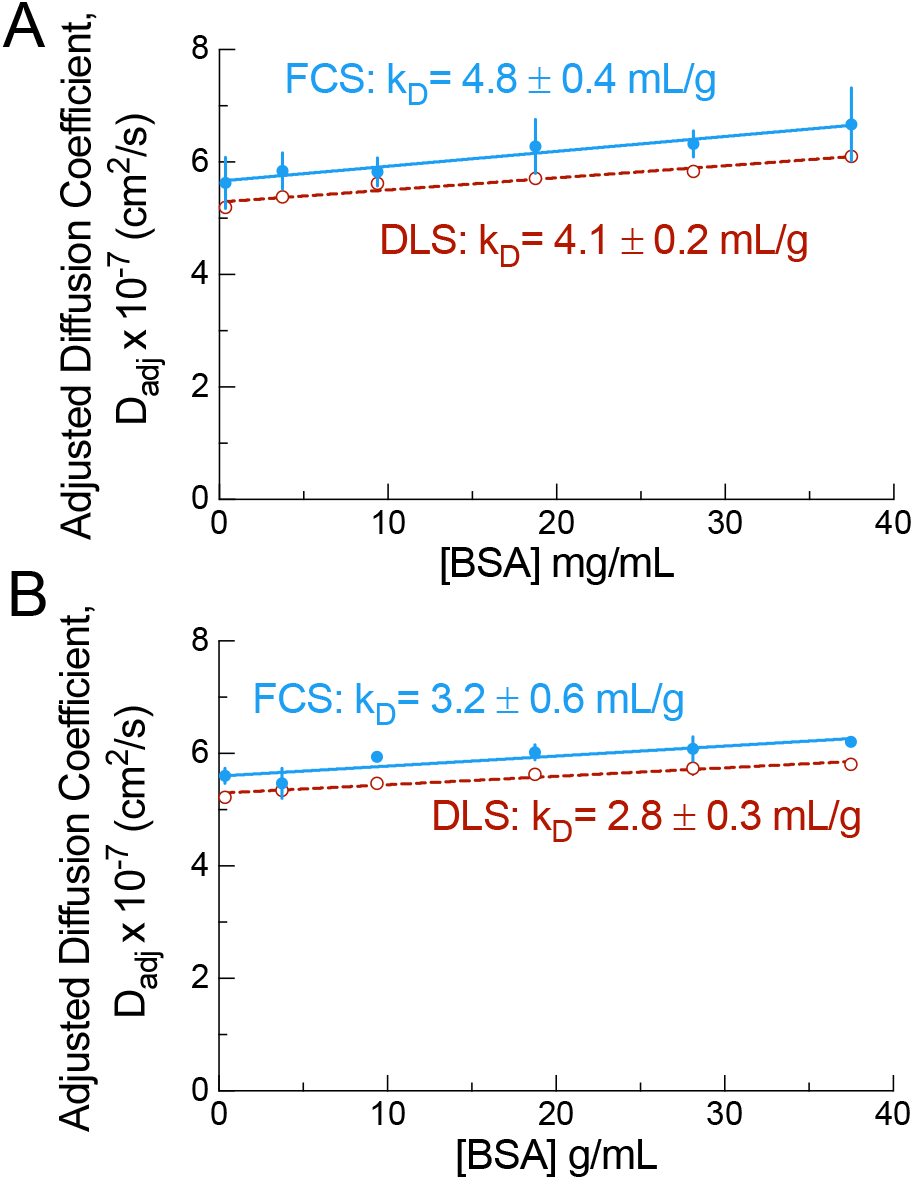
Comparison of diffusion coefficients determined by FCS and DLS for BSA at pH 7.4 (A) and pH 6.0 (B). The second virial coefficient, B_22_M (or k_D_) for DLS measurements were obtained using the average diffusion coefficient from a complete distribution. This had only a minor effect on kD measured at pH 7.4 (4.1 ± 0.2 mL/g) compared to the value (4.3 ± 0.4 mL/g) obtained from the average diffusion coefficient of the monomeric peak and reported in Table 1. Using the complete diffusion coefficient distribution also had a minor effect on the DLS kD value measured at pH 6.0 (2.8 ± 0.3 mL/g) compared to our reported value (3.4 ± 0.3 mL/g). In both cases, the average diffusion coefficient of the monomeric peak resulted in better agreement between FCS and DLS virial coefficients, as displayed in Fig. 1 of the Results section. However, DLS values are still comparable to FCS values when using the average diffusion coefficient of the complete distribution.

## AUTHOR CONTRIBUTIONS

H.A.L., A.N., and W.M.A. designed experiments. H.A.L. performed experiments and analyzed data. A.N. and W.M.A. provided analytical tools. H.A.L., A.N, and W.M.A wrote the manuscript.

## ACKNOWLEDGEMENTS

This work was supported by the NIH (T32GM007750) and the UWSOP Faculty Innovation Fund. The authors gratefully acknowledge Genentech and Dr. Mark Chiu (Janssen Pharmaceuticals) for providing IgG samples. We thank John Sumida for completing buffer density and viscosity measurements. We also thank Dr. John Correia for constructive feedback.

## Notes

### Competing Interest Statement

The authors have declared no competing interest.

